# GDF3 is an obesity-induced regulator of TGFβ superfamily signaling

**DOI:** 10.1101/2022.11.07.515236

**Authors:** Deepti Ramachandran, Nagasuryaprasad Kotikalapudi, Gregory R. Gipson, Luca Troncone, Kylie Vestal, David E. Maridas, Anton Gulko, Linus T. Tsai, Vicki Rosen, Paul Yu, Thomas B. Thompson, Alexander S. Banks

## Abstract

Growth differentiation factor 3 (GDF3) is a relatively understudied member of the TGFβ superfamily that is highly expressed during development. However, the function of GDF3 in adult biology is contentious. We use *in vivo* approaches to show that GDF3 loss-of-function in adipose tissue of obese adult mice causes reduced body weight and improved whole-body insulin sensitivity. These effects are accompanied by altered regulation of genes targeted by the TGFβ superfamily *in vivo*. Using *in vitro* approaches, we show that GDF3 can influence both arms of the TGFβ superfamily: GDF3 simultaneously inhibits BMP signaling and activates activin-like SMAD 2/3 signaling. We identify the type II receptors mediating this activity. GDF3 binds to the type II receptors BMPR2, ACTRIIA and ACTRIIB and achieves dose-dependent inhibition of multiple BMP proteins including BMP2, BMP7, BMP9, BMP10, and BMP15 *in vitro*. We also find that GDF3 activates TGFβ/activin-like SMAD2/3 signaling. Unbiased expression profiling confirms that GDF3 both attenuates BMP2-regulated gene expression and drives TGFβ/activin-like gene expression. Together these results provide much needed clarity to both the molecular pathways involved in GDF3 signaling and the physiological effects of GDF3 loss of function.

## INTRODUCTION

GDF3 is a member of the TGFβ superfamily. Proteins in this family are translated into full-length pre-proteins that undergo enzymatic cleavage to produce mature peptides (*1*). The mature GDF3 polypeptide lacks a highly conserved cysteine found in nearly all BMPs/GDFs and two additional cysteines found in TGFβs and activins (*2, 3*). Phylogenetic analysis of the 114 amino acid mature human GDF3 protein aligned to mature polypeptides of the TGFβ superfamily shows closest amino acid identity to GDF1 at 48% (*2*). The GDF3/GDF1 phylogenetic branch maps marginally closer to the mature TGFβ, activin, GDF15, GDF11, and myostatin/ GDF8 protein branches. The mature GDF3 protein clustered marginally more distantly from BMPs and other GDF proteins (Fig 1A).

**Figure 1:**
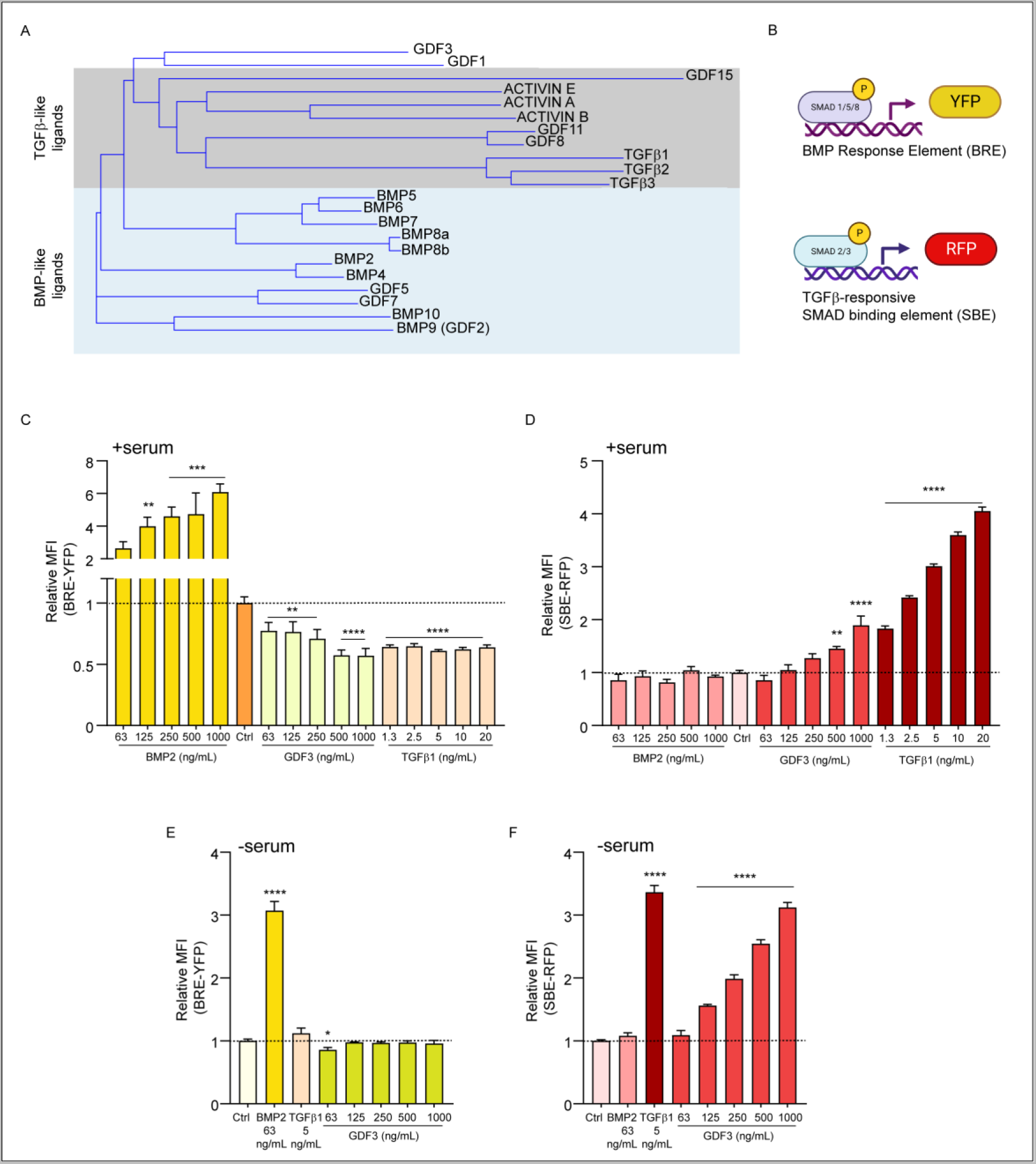
GDF3 simultaneously inhibits BMP signaling and activates TGFβ/activin-like signaling. A) Phylogenetic analysis following multiple sequence alignment of mature human GDF3 protein and mature polypeptides of members of the TGFβ superfamily. B) Schematic representations of the BMP responsive element (BRE) driving yellow fluorescent protein (YFP) and a TGFβ/activin responsive SMAD binding element {SBE) driving red fluorescent protein (RFP). C-G) C2C12 myoblasts stably expressing both the BRE-YFP and SBE-RFP reporters (dual reporter cells), C) Relative Mean Fluorescence Intensity (MFI) of YFP in dual reporter cells incubated with increasing doses of recombinant BMP2, GDF3, and TGFβl in complete media containing 20% fetal bovine serum for 24 hours. D) Relative MFI of RFP in the same cells as C. E) Relative MFI of YFP in reporter cells incubated with recombinant proteins in serum-free media for 24 hours F) Relative MFI of RFP in the same cells as E. *p<0.05, **p<0.01, ***p<0.001, ****p<0.0001 compared to untreated control (Ctrl),

Significant aspects of the biology of the protein Growth Differentiation Factor 3 (GDF3) are either unknown or disputed. Like other members of the TGFβ superfamily, GDF3’s function may be highly cell type and context-dependent. It is highly expressed in stem cell populations and helps to maintain markers of pluripotency, however, Gdf3 mRNA levels decline sharply after differentiation (*4, 5*). GDF3’s function in differentiated cells is less clear (*6, 7*). In adult humans and mice, Gdf3 mRNA levels sharply increase in obesity, aging, inflammation, and following an ischemic event (*8-12*). Understanding whether GDF3 has a role in regulating these diseases requires a mechanistic understanding of GDF3 biology.

The nuclear hormone receptor PPARγ controls Gdf3 expression (*10*). Obesity increases GDF3 levels, while GDF3 expression is reduced by tissue-specific PPARγ knockout, treatment with the insulin-sensitizing PPARγ ligands (thiazolidinediones), or by mutating PPARγ such that it cannot be phosphorylated at the 273^rd^ amino acid (Serine S273, a post translational modification that positively correlates with increasing weight gain and obesity) (*9, 13-15*). In all these examples, high GDF3 levels positively correlate with insulin resistance, and decreased GDF3 levels correlate with insulin sensitivity. Moreover, exogenously increasing GDF3 expression with adeno-associated virus (AAV) was sufficient to make even lean mice moderately insulin resistant (*9*). Together, these findings suggest that increased GDF3 levels may contribute to obesity-induced insulin resistance, positioning GDF3 as a candidate target gene to develop therapies for type 2 diabetes, obesity, or other indications.

Conflicting research exists in the field about GDF3’s mechanism – some studies suggest that GDF3 inhibits BMP signaling while others maintain that GDF3 activates TGFβ signaling (*4, 9, 11, 16-19*).

To study the molecular function of GDF3, we created tools to interrogate GDF3’s role in BMP and TGFβ signaling simultaneously. We developed cells expressing dual fluorescent reporters, and showed that GDF3 can simultaneously inhibit BMP signaling and stimulate activin-like signaling. We identified three different type II receptors of the TGFβ family that bind GDF3: BMPR2, ACTRIIA and ACTRIIB. We also found that both recombinant and genetically encoded GDF3 induce a gene expression signature similar to TGFβ1 while blocking BMP signaling. We examine an inducible model of Gdf3 deletion in adult male obese mice to demonstrate the loss of function effects of GDF3 in obese adipose tissue on gene expression and physiology. These studies suggest that GDF3 has dual roles to antagonize BMP signaling and to promote activin-like or myostatin-like signaling, and that reducing GDF3 levels in obesity could improve metabolic health.

## RESULTS

### Generation of cells expressing a dual fluorescence reporter for simultaneous BMP (BRE) and TGFβ (SBE) signaling

BMP signaling promotes phosphorylation of SMAD1, SMAD5, and/or SMAD8. Similarly, TGFβ & activin signaling promote the phosphorylation of SMAD2, and/or SMAD3. Immunoblotting for protein phosphorylation has served as the backbone for understanding ligand and receptor signaling (*20*). Luminescence reporter assays for BMP signaling have used the Id1 promoter’s <u>B</u>MP-<u>R</u>esponsive <u>E</u>lement (BRE) (*21*). Likewise, TGFβ signaling has used luminescence driven by the <u>S</u>MAD3-<u>B</u>inding <u>E</u>lement (SBE or CAGA_12_) (*22*). The Elowitz group at Caltech recently created a flow cytometry-compatible BMP reporter with the BRE promoter driving expression of Citrine, a yellow fluorescent protein (referred to as BRE-YFP henceforth) (*23-25*). This has further increased throughput and allowed for a systems-level approach to BMP signaling not previously possible. We extended these studies to generate a red fluorescent TGFβ signaling reporter driven by the SBE promoter (referred to as SBE-RFP henceforth). We previously showed that *Gdf3* is expressed in adipose tissue monocytes, macrophages, and adipocytes during obesity. In addition, lower *Gdf3* levels in adipose tissue correlate with improved insulin sensitivity and glucose uptake in skeletal muscle tissue (*9*). We therefore used the C2C12 muscle cell line to study the molecular function of GDF3. In these cells, Gdf3 mRNA levels were undetectable by RNA-seq or qPCR making them an ideal model for gain-of-function studies. C2C12 myoblasts were transfected with both the BMP responsive BRE-YFP reporter construct and the TGFβ responsive SBE-RFP construct (Fig 1B) to generate C2C12 myoblasts stably expressing both constructs (dual reporter cells). To prevent cellular differentiation, our initial experiments were performed in complete C2C12 media containing 20% serum. We found, as previously published, recombinant BMP2 potently activated the BRE-YFP reporter in a dose-dependent manner (Fig 1C) (*23*). Similarly, we demonstrated that TGFβ1 potently activated the SBE-RFP reporter in a dose-dependent manner (Fig 1D). While BMP2 had no effect on the SBE-RFP reporter at any dose tested, all the tested doses of TGFβ1 inhibited the BRE-YFP reporter by ∼30%. We conclude that the fluorescent dual reporter system faithfully recapitulates the results from prior luciferase based assays and now allows us to examine the effect of TGFβ superfamily ligands on both BMP and TGFβ reporters simultaneously in the same cells.

### GDF3 simultaneously inhibits BMP signaling and activates TGFβ signaling in a dose dependent manner

We next sought to understand the effects of GDF3 in this system. Dual reporter cells were incubated with increasing doses of recombinant mouse GDF3 in complete media. We observed recombinant GDF3 could inhibit the BRE-YFP reporter in a dose-dependent manner (Fig. 1C). GDF3, like TGFβ1, activated the SBE-RFP reporter but required much higher concentrations than TGFβ1; 1000 ng/mL of GDF3 had a similar effect as 1.3 ng/mL of TGFβ1 (Fig. 1D). Considering serum contains a mixture of BMP and TGFβ ligands that could influence signaling, we next decided to test the effects of our recombinant proteins under serum-free conditions.

To do this, the dual reporter cells were incubated in serum-free media for 4 hours, followed by treatment with recombinant proteins for 24 hours. Under serum-free conditions, recombinant GDF3 had a minimal effect on the BRE-YFP reporter while the strong reporter activation of BMP2 was further enhanced in the absence of serum (Fig. 1E). In contrast, recombinant GDF3 activated the SBE-RFP reporter more potently in serum-free conditions (Fig 1F). TGFβ1 also strongly activated the SBE-RFP reporter (Fig 1F). These data suggest that while GDF3 can autonomously activate TGFβ signaling, it requires a serum-derived factor to affect BMP signaling.

### GDF3 is a BMP signaling antagonist with a high affinity for the receptor BMPRII

Considering the different effects of GDF3 on BMP signaling in the presence or absence of serum, we hypothesized that GDF3 may inhibit BMP signaling only in the presence of BMP ligands. We therefore tested the effect of recombinant GDF3 on isolated recombinant BMP2, BMP7, BMP9, BMP10, or BMP15 in serum-free conditions. Indeed, while GDF3 alone had no effect on the BRE-YFP reporter, GDF3 inhibited all BMPs tested in a dose dependent manner (Fig. 2A).

**Figure 2:**
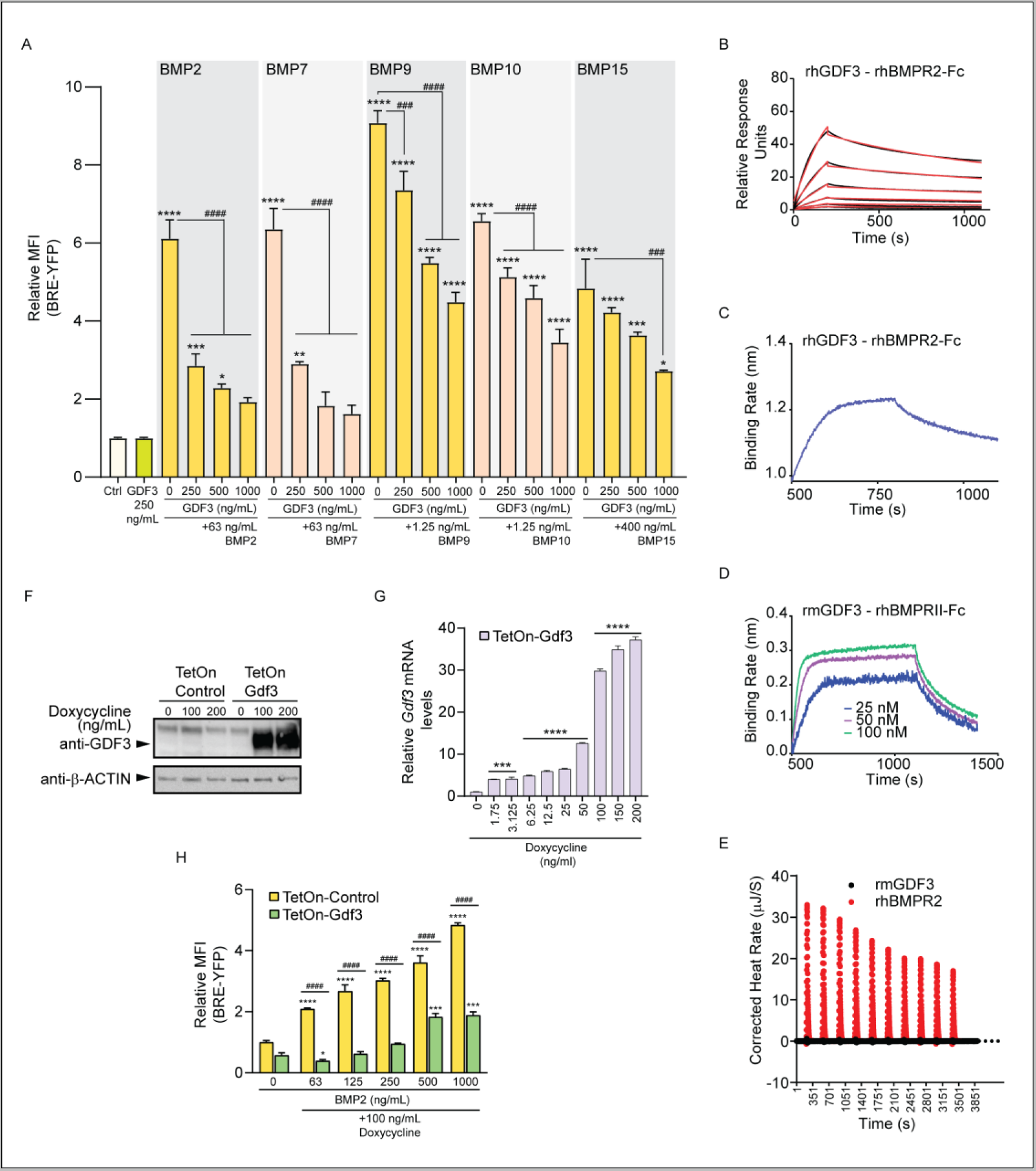
GDF3 is a BMP signaling antagonist with a high affinity for the receptor BMPRII. A) Relative MFI of YFP in C2C12 dual reporter cells incubated with recombinant human (rh) BMP2, recombinant mouse (rm) BMP7, rhBMP9, rmBMPIQ or rhBMP15 and/ or increasing doses of rmGDF3 in serum free media for 24 hours. B) Surface Plasmon Resonance (SPR) analysis of rh-GDF3 with Fc chimera of rhBMPRII. The raw binding curves are in black and the kinetic binding fits are in red. C) Biolayer Interferometry (BLI) analysis of 50nM rhGDF3 interacting with rhBMPRII-Fc. D) BLI analysis of rmGDF3 with rhBMPRII-Fc. E) Binding isotherm for the titration of GDF3 against rhBMPRII-Fc. F) Western blot analysis of 293T cells transiently transfected with doxycycline inducible control (TetOπ-Control) or mouse Gdf3 (TetOn-Gdf3) vectors for 24 hours followed by doxycycline treatment for 48 hours. G-H) C2C12 dual reporter cells were transiently transfected with TetOn-Control or TetOn-Gdf3 vectors for 24 hours followed by doxycycline treatment in serum free media for 48 hours. All cells were either harvested for RNA isolation (G) or cells expressing YFP, RFP and BFP were sorted via flow cytometry and analyzed for changes in reporter activity (H). G) Relative mRNA fold change of Gdf3 mRNA in response to increasing doses of Doxycycline H). Relative MFI of YFP in sorted YFP, RFP and BFP positive cells treated with rhBMP2 for the last 24 hours. */#p<0.05, *·/##p<0.01, ***/###p<0.001, ****/#### p<O.0001 when compared to control (*) or BMP ligand alone (*).

These findings suggest that GDF3 is a broad inhibitor of BMP signaling. To determine if GDF3 acts as a pan BMP inhibitor via BMP signaling receptor interaction, we performed ligand receptor interaction assays using three different techniques. Surface Plasmon Resonance (SPR) analysis of recombinant human GDF3 with the Fc-chimera of recombinant human type II BMP receptor BMPRII showed that GDF3 has a high affinity for BMPRII (Fig 2B and Table 1). This interaction was also confirmed with Biolayer Interferometry (BLI) analysis which showed that both human and mouse recombinant GDF3 could bind to BMPRII with a high affinity (Fig 2C and 2D). Isothermal Titration Calorimetry (ITC) with recombinant mouse GDF3 against BMPRII also showed interaction between this ligand and receptor combination (Fig 2E and Table 1). These three complementary assays all confirm a strong binding interaction of GDF3 to BMPR2.

**Table 1:**
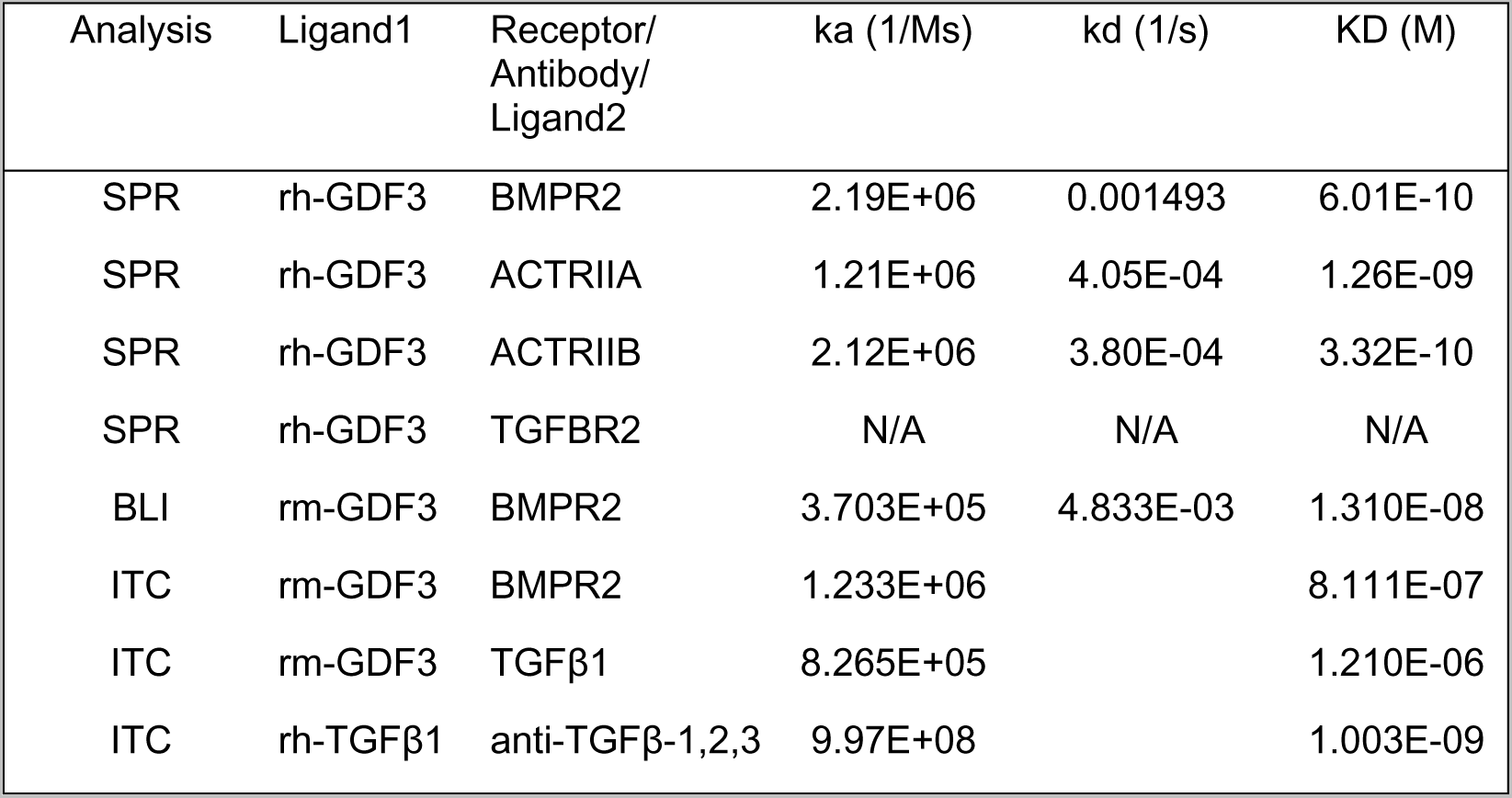
Kinetics data from protein - protein interaction studies.

| Analysis | Ligand1 | Receptor/<br>Antibody/<br>Ligand2 | ka (1/Ms) | kd (1/s) | KD (M) |
| --- | --- | --- | --- | --- | --- |
| SPR | rh-GDF3 | BMPR2 | 2.19E+06 | 0.001493 | 6.01E-10 |
| SPR | rh-GDF3 | ACTRIIA | 1.21E+06 | 4.05E-04 | 1.26E-09 |
| SPR | rh-GDF3 | ACTRIIB | 2.12E+06 | 3.80E-04 | 3.32E-10 |
| SPR | rh-GDF3 | TGFBR2 | N/A | N/A | N/A |
| BLI | rm-GDF3 | BMPR2 | 3.703E+05 | 4.833E-03 | 1.310E-08 |
| ITC | rm-GDF3 | BMPR2 | 1.233E+06 |  | 8.111E-07 |
| ITC | rm-GDF3 | TGFβ1 | 8.265E+05 |  | 1.210E-06 |
| ITC | rh-TGFβ1 | anti-TGFβ-1,2,3 | 9.97E+08 |  | 1.003E-09 |

### Genetically encoded Gdf3 and recombinant GDF3 both inhibit BMP/BRE reporter activity

Noting previous concerns about TGFβ contamination in purified recombinant proteins derived from CHO cells (*26*), we wanted to ensure that our effects were not due to trace levels of TGFβ protein in our recombinant GDF3 preparations. We designed vectors using the Tet-On inducible gene expression system such that we could temporally control the expression of a control protein, blue fluorescent protein (BFP) alone (TetOn-Control), or full length mouse GDF3 and BFP (TetOn-Gdf3) in the presence of the tetracycline analog doxycycline. Western blot analysis of cells transfected with the TetOn-Control or TetOn-Gdf3 vectors showed strong full-length GDF3 protein expression only in TetOn-Gdf3 cells and only in the presence of doxycycline (Fig 2F). We next transfected our dual reporter cells with either the control or GDF3 encoding vectors and treated these cells with increasing doses of doxycycline under serum free conditions for 48 hours. Doxycycline treatment led to a significant dose-dependent increase in Gdf3 mRNA in TetOn-Gdf3 expressing cells (Fig 2G). Gdf3 mRNA remained undetectable in cells expressing only BFP at any dose of doxycycline tested (data not shown). We next tested if the induction of genetically encoded Gdf3 could activate the BRE-YFP reporter. Doxycycline induction of Gdf3 showed no changes in BRE-YFP reporter activity (Fig S1A), consistent with results from recombinant GDF3 in serum free conditions (Fig 1E). Doxycycline induced GDF3 however, was able to attenuate the response of the BRE-YFP reporter to BMP2 signaling (Fig 2H). These results are consistent with our reporter assays using recombinant GDF3 and BMP2 (Fig 2A). Doxycycline induction of control cells had no effect on the BRE-YFP reporter (Fig S1B).

### GDF3 acts as a TGFβ/activin-like SMAD2/3 signaling agonist

In contrast to GDF3’s inability to affect BMP signaling under serum free conditions, recombinant GDF3 activated the SBE-RFP more strongly in the absence of serum in the media (Fig 1F). TGFβ1 also potently activated the SBE-RFP reporter (Fig 1F). These data suggest that GDF3 can autonomously activate SMAD2/3 signaling. To identify which receptors might be involved in GDF3’s ability to activate the SBE-RFP reporter we performed ligand - receptor interaction studies with recombinant GDF3 and type II receptors in the TGFβ signaling pathway. SPR analysis of recombinant human GDF3 with recombinant human ACTRIIA, ACTRIIB and TGFBR2 showed that GDF3 had a high affinity for ACTRIIA and ACTRIIB, but had no binding affinity for TGFBR2 (Fig 3A and Table 1). Surprisingly, parallel BLI analysis with recombinant human and mouse GDF3 found high ligand affinity for ACTRIIA but no affinity for ACTRIIB or TGFBR2 (Fig 3B). We also investigated how GDF3 would interact with TGFβ-like signaling ligands to affect SBE-RFP reporter activity. To do so, we added recombinant mouse GDF3 with either recombinant TGFβ1 or activin A in serum-free conditions. We found that GDF3 had additive effects on SBE-RFP reporter activity with both TGFβ1 and activin A. GDF3 in combination with TGFβ1 or activin A leads to significantly higher SBE-RFP reporter activity than either ligand in isolation (Fig 3C and 3D). These results suggest that GDF3 acts as a SMAD2/3 signaling ligand in the absence of serum or in the presence of other SMAD2/3 signaling ligands.

**Figure 3:**
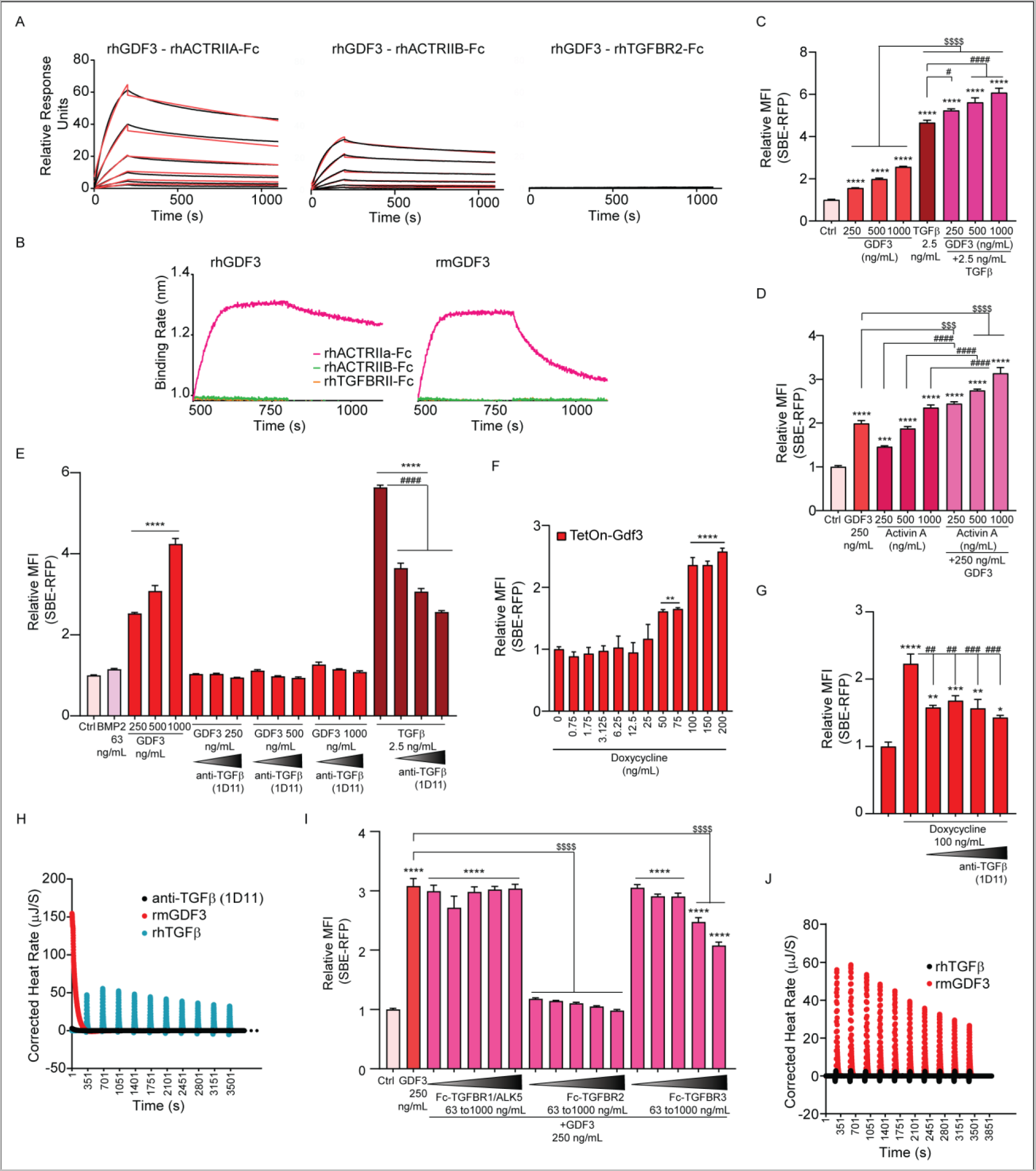
GDF3 acts as a TGFβ/ activin like SMAD2/3 signaling agonist. A) SPR analysis rh-GDF3 with rh-ACTRIIA-Fc, rh-ACTRIIB-Fc and rh-TGFBR2-Fc. B) BLI analysis of 50nM rhGDF3 or rmGDF3 with rh-ACTRIIA-Fc, rh-ACT-RIIB-Fc and rh-TGFBR2-Fc. C) Relative MFI of RFP in C2C12 dual reporter cells incubated with recombinant rmGDF3 and/or rh-TGFβl. D) Relative MFI of RFP in cells incubated with rm-GDF3 and/or rh-activinA. E) Relative MFI of RFP in cells incubated with rh-BMP2, rm-GDF3 or rh-TGFβ1 with or without the pan-TGF β specific antibody (Clone 1D11). F-G) C2C12 dual reporter cells were transiently transfected with the TetOn-Gdf3 plasmid followed by doxycycline treatment in serum free media for 48 hours (F). Anti-TGFβ (Clone 1D11) antibody was added to the cells for the last 24 hours (G). Cells expressing YFP, RFP and BFP were sorted via flow cytometry and analyzed for changes in RFP reporter activity (F, G). H) Binding isotherm for the titration of rm-GDF3 or rh-TGFβ1 against the anti-TGFβ (1D11) antibody. I) Relative MFI of RFP in cells incubated with recombinant Fc chimeras of the receptors TGFBR1/ALK5, TGFBR2 and TGFBR3 and/or recombinant GDF3. J) Binding isotherm for the titration of rmGDF3 against rhTGFβl. */ $ /#p<0.05, **/$$/##p<0.01, ***/$$$/###p<0.001, ****/$$$$/#### p<0.0001 when compared to control (*) or GDF3 alone ($) or the BMP or TGFβ signaling ligand (#)

### Genetically encoded Gdf3 and recombinant GDF3 both increase activin/SBE reporter activity

To address the issue of trace levels of contaminating TGFβ in CHO cell derived recombinant protein samples (*26*), we added the pan TGFβ neutralizing antibody (clone 1D11) to our reporter assay and saw that in the presence of the 1D11 antibody, the ability of recombinant GDF3 to activate the SBE-RFP reporter was completely abrogated (Fig 3E). This indicated that GDF3’s activation of TGFβ signaling might be due to trace levels of TGFβ contamination. To address this concern, we used our inducible genetically encoded TetOn-Gdf3 or control vectors to eliminate the contribution of possible contaminants. We saw that doxycycline induction of Gdf3 led to a dose dependent increase in the SBE-RFP reporter activity (Fig 3F), indicating that GDF3 is indeed able to activate the SBE-RFP reporter. Doxycycline induction of BFP in dual reporter cells did not lead to any effect on the SBE-RFP reporter (Fig S1C). Surprisingly, we also showed that the TGFβ neutralizing antibody (1D11) could blunt the activation of the SBE-RFP reporter seen in response to doxycycline induction of genetically encoded Gdf3 (Fig 3G). To rule out the possibility that the 1D11 antibody might be binding and neutralizing GDF3 directly, we performed ITC assays with the 1D11 antibody and recombinant mouse GDF3 or human TGFβ1. While recombinant TGFβ1 was able to generate heat when titrated against the TGFβ neutralizing 1D11 antibody indicating an interaction between these two molecules, recombinant GDF3 showed no interaction with the antibody (Fig 3H). To further control for the specificity of this interaction, we also performed a BLI analysis with the TGFβ neutralizing antibody and recombinant GDF3, BMP9 or TGFβ3. While recombinant human TGFβ3 showed high affinity for the 1D11 antibody, recombinant human and mouse GDF3 as well as human BMP9 showed very low affinity for the antibody (Fig S1D). These results suggest that GDF3’s activation of TGFβ signaling might require the presence of latent TGFβ in the cells either through a direct or indirect interaction between GDF3 and TGFβ. We next investigated whether GDF3’s induction of the SBE-RFP reporter could be blocked by soluble forms (Fc chimeras) of the canonical TGFβ receptors TGFBR1 (ALK5), TGFBR2 or the TGFβ co-receptor TGFBR3 (Betaglycan). ALK5-Fc had no effect on GDF3’s induction of the SBE-RFP reporter (Fig 3I). The TGFBR3-Fc was able to block GDF3’s effect to a small extent at the highest concentrations tested. However, TGFBR2 was able to completely block the effect of recombinant mouse GDF3 on the SBE-RFP reporter even at the lowest concentration tested (Fig 3I). Together with our SPR and BLI data which show that GDF3 does not bind to TGFBR2, these data indicate that GDF3 might be activating the SBE-RFP reporter either through direct or indirect interaction with TGFβ. Indeed, ITC analysis of recombinant mouse GDF3 titrated against recombinant human TGFβ1 demonstrated that GDF3 interacts with TGFβ1 (Fig 3J). These data place GDF3 as a unique SMAD2/3 signaling molecule that potentially interacts with latent TGFβ to activate TGFβ signaling.

### GDF3 promotes a gene expression profile associated with decreased muscle cell differentiation

We next sought to validate the effects of GDF3 on endogenous gene expression by performing RNA-sequencing experiments in our dual reporter cells. Recombinant mouse GDF3 increased expression of 92 genes (G+) and decreased expression of 126 genes (G-). While TGFβ1 had more pronounced effects to increase expression of more than 1000 genes (T+) and to repress the expression of greater than 1300 genes (T-) beyond the significance threshold. Approximately 44% of GDF3 regulated genes were also significantly co-regulated by TGFβ1 (Fig 4A). Of note, GDF3 at 250 ng/ml had effects to less strongly regulate gene expression when compared to 2.5 ng/ml TGFβ1 (Fig 4A). This weaker effect of Gdf3 on endogenous gene expression compared to TGFβ1 is consistent with results from our reporter assay experiments (Fig 1F).

**Figure 4:**
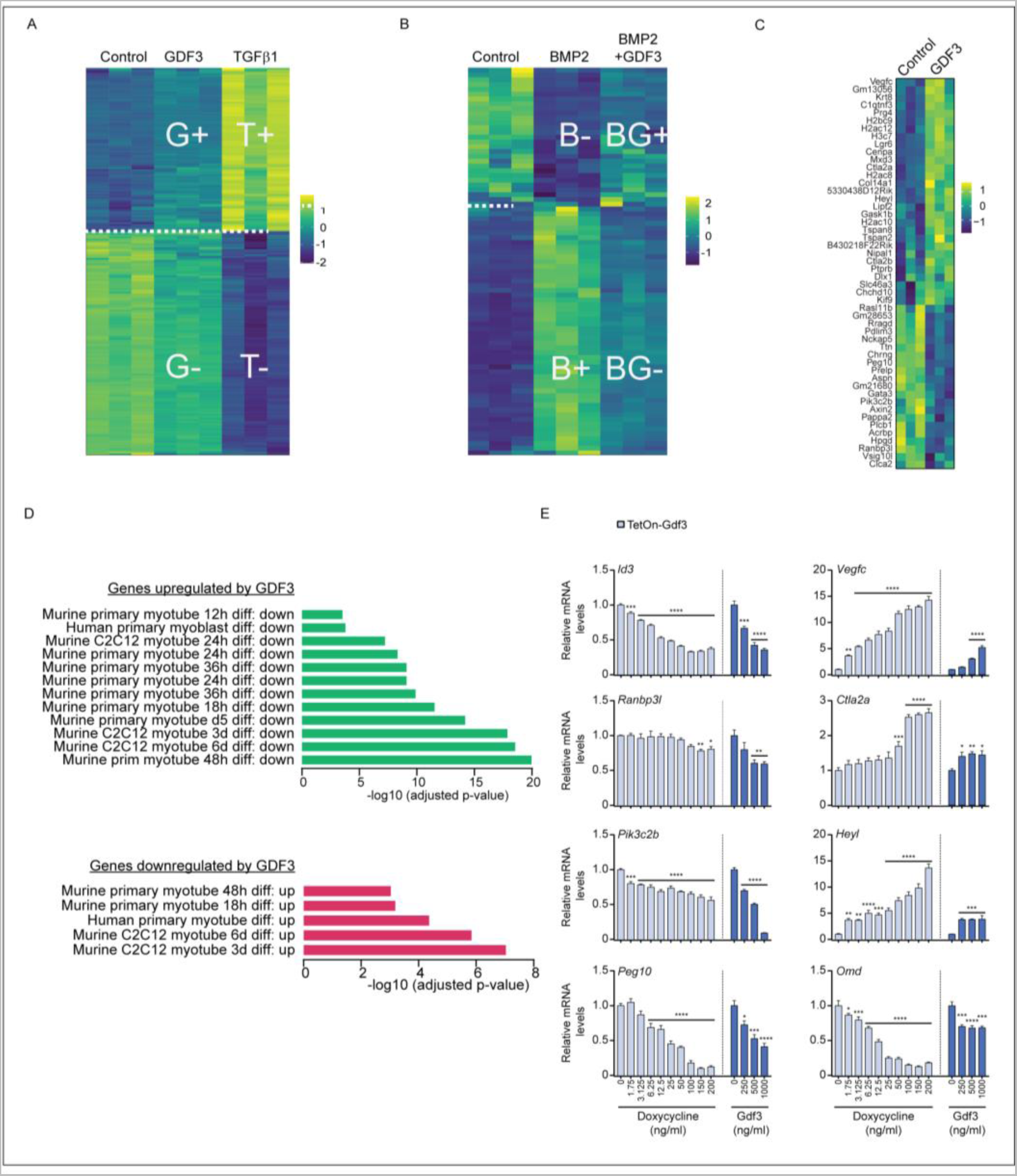
GDF3 inhibits C2C12 myoblast differentiation. A-D) RNA sequencing data from C2C12 dual reporter cells incubated in serum free media (Control) or with recombinant proteins in serum free media for 24 hours. A) Heat map of gene expression changes from cells incubated with Recombinant GDF3 or TGFβl. B) Heat map of gene expression chang­es from cells incubated with recombinant BMP2 or BMP2 and GDF3. C) Heat map for selected gene expression changes in cells from (A) treated with recombinant GDF3 versus Control. D) KEGG pathway analysis of RNA sequencing data from A and B. E) C2C12 dual reporter cells were transiently transfected with a plasmid encoding doxycycline inducible mouse Gdf3 and blue fluorescent protein (TetOn-Gdf3) followed by doxycycline treatment in serum free media for 48 hours or incubated with recombinant GDF3 for 24 hours.

Recombinant BMP2 is a potent regulator of transcriptional activity with more than 580 genes significantly upregulated (B+). We found that GDF3 significantly limited the expression of 47 genes activated by BMP2 (BG+). Similarly, of the more than 570 genes repressed by BMP2 (B-), the decreased expression of 66 genes was blocked with the addition of GDF3 (BG-) (Fig 4B). GDF3 limited the regulation of approximately 10% of the genes controlled by BMP2, suggesting that GDF3 can act as a limited antagonist of BMP2 activity on endogenous gene expression. In our experiments, cells treated with GDF3 had strong downregulation of *Ranbp3l*, a gene implicated in SMAD signaling (Fig 5C). Other notable downregulated genes included *Pik3c2b, Id3, Omd,* and *Peg10*. Among strongly upregulated genes are *Ctla2a*, *Heyl*, and *Vegfc* (Fig 4C).

**Figure 5:**
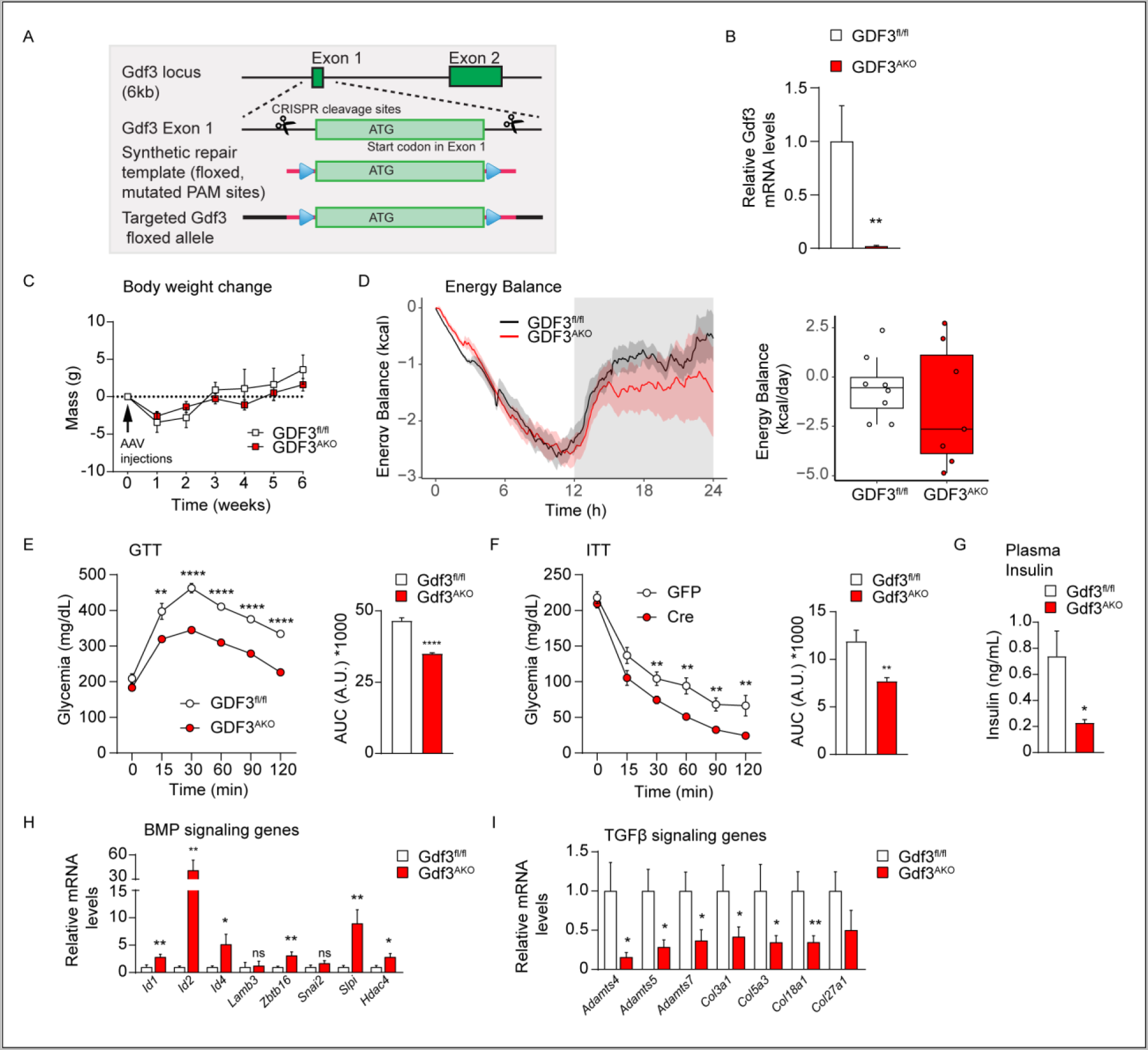
GDF3 loss of function in epididymal adipose tissue leads to improved metabolic health under conditions of diet induced obesity. A) Schematic representation of floxed GDF3 locus used to generate GDF3^fl,fl^ mice using CRISPR/Cas9 technology. B-l) Gdf3^fl/fl^mice fed high fat diet for 8 weeks were injected with either control AAV-eGFP (GDF3^fl,fl^) or AAV-Cre (Gdf3^AKO^) in their eWAT (n = 8/9). Week of AAV injection (week 0). B) RT-qPCR analysis of Gdf3 mRNA levels in the eWAT of these mice in week 6. C) Body weight changes from AAV injection (week 0) until sacrifice (week 6). D) Energy balance analysis over 24h (left), average daily energy balance (right) in week 3. E) Glucose tolerance test (GTT) in week 2 (glucose dose: 1 g/kg BW, intraperitoneal injections) with area under the curve (AUC) on the right. F) Insulin tolerance test (ITT) in week 5 (Insulin dose: 0.75 U/kg BW) with AUC on the right. G) Plasma insulin levels after 4 h fast,in week 5. H-l) RT-qPCR analysis of BMP signaling target genes (H) and TGFβ signaling target genes (I) in eWAT in week 6. *ρ<0.05, **p<0.01, **‘p<0.001, ****p<0.0001 compared to control (Gdf3^flΛ1^)

The transcriptional profile of cells treated with GDF3 is consistent with inhibition of muscle cell differentiation. That is, the genes downregulated in GDF3-treated cells overlap with genes upregulated during the differentiation of murine primary myoblasts, murine C2C12 myoblasts, or human myoblasts into myotubes (*27, 28*). Conversely, genes upregulated following GDF3 treatment overlap with the genes that are inhibited during myoblast to myotube differentiation (Fig 4D) (*29*). These results indicate that GDF3 might inhibit differentiation of C2C12 myoblasts to myotubes.

To determine if the gene expression changes seen with recombinant GDF3 hold true with genetically encoded Gdf3, we analyzed the expression levels of selected target genes regulated by recombinant GDF3 using RT-qPCR analysis. The expression levels of all the genes tested were regulated similarly in the presence of increasing doses of recombinant GDF3 or alternately, in the presence of increasing doses of doxycycline in cells expressing the TetOn-Gdf3 vector (Fig 4E). Of these, *Id3*, *Ranbp3l*, *Pik3c2b*, *Peg10,* and *Omd* all showed significantly lower gene expression levels in response to induction of genetically encoded Gdf3 or recombinant GDF3, while *Vegfc*, *Ctla2a* and *Heyl* showed increased gene expression levels in response to genetically encoded Gdf3 or recombinant GDF3 (Fig 4E). These experiments find similar regulation of endogenous gene expression by recombinant GDF3 and a genetically encoded *Gdf3*, assuaging concerns that the effects of recombinant GDF3 could be an effect of trace amounts of TGFβ. Together these results suggest that GDF3 does activate SMAD2/3 signaling, albeit in combination with latent TGFβ.

### Gdf3 loss of function in adipose tissue of diet induced obese mice leads to weight loss and improved metabolic health

We next sought to understand the biology of Gdf3 in developmentally mature mice. To study the impact of loss of function of Gdf3 in adulthood, we generated mice bearing a conditional allele of Gdf3, such that exon 1, encoding the Gdf3 start codon, is flanked by two loxP sites. Gdf3 is conditionally deleted in the presence of Cre recombinase (Figure 5A). Considering the high expression of GDF3 in adult adipose tissue during obesity (*9, 11, 30, 31*), we fed Gdf3^fl/fl^ mice high fat diet for 8 weeks, followed by epididymal white adipose tissue directed administration of either a control adeno associated virus (AAV-GFP), or AAV-Cre to generate eWAT-specific Gdf3 knock out (Gdf3^AKO^) mice as seen by reduced Gdf3 mRNA in the eWAT of Gdf3^AKO^ mice (Fig 5B). We found no difference in the body weight gain between Gdf3^fl/fl^ mice and Gdf3^AKO^ mice after AAV administration (Fig 5C) but Gdf3^AKO^ mice were in negative energy balance as measured by indirect calorimetry (Fig 5D) and had significantly lower body weights than Gdf3^fl/fl^ mice beginning from 3 weeks post AAV administration (Fig S1E). Consistent with this lower body weight, Gdf3^AKO^ mice showed significantly improved glucose tolerance and improved insulin sensitivity during a Glucose Tolerance Test (GTT) and Insulin Tolerance Test (ITT) respectively (Fig 5E and 5F). The AAV-Cre mice also had reduced fasting insulin levels compared to controls (Fig 5G). Gene expression in knockout epididymal adipose tissue revealed increased expression of genes downstream of BMP signaling including *Id1*, *Id2*, *Id4*, *Zbtblb*, *Slpi* and *Hdac4* (Fig 5H). Gdf3^AKO^ mice also showed reduced expression of TGFβ signaling target genes including *Adamts4*, *Adamts5*, *Adamts7*, *Col3a1*, *Col5a3*, *Col18a1* and *Col27a1* (Fig 5I). These data support the hypothesis that Gdf3 regulates both BMP signaling as well as TGFβ signaling in adulthood.

## DISCUSSION

### GDF3, a bi-functional protein

Our understanding of the molecular role of GDF3 has evolved beyond the unilateral control of either BMP or TGFβ signaling into that of a dual regulator (Fig 6A and 6B). GDF3’s role as a BMP inhibitor was first demonstrated in human embryonic stem cells, frog embryos, and mouse gene-trap models (*4*). These studies illustrated that both recombinant GDF3 and exogenous Gdf3 cDNA can inhibit BMP reporter activity. Contemporaneously, others found that GDF3 acted as a Nodal-like ligand activating the SMAD 2/3 pathway similarly to TGFβ in the pre-gastrulation mouse embryo (*18*). These findings led the authors to the “intriguing possibility that GDF3 acts as a bi-functional protein” with the caveat that while endogenous levels of GDF3 could inhibit BMP signaling, only extremely high, non-physiological levels of GDF3 acted as a TGFβ signaling activator (*3, 19*). Our results support the assertion that GDF3 is a bi-functional protein at the tested concentrations. One caveat to our work and to that of others is that we lack an accurate estimation of endogenous GDF3 levels due to the challenges of antibody based detection systems within the TGFβ-superfamily (*32*). Regardless, we find that GDF3 has similar potency to recombinant activin A in our reporter system (Fig 3D). We also demonstrate that the same concentrations of recombinant GDF3 are sufficient to both inhibit BMP mediated SMAD 1/5/8 signaling and activate TGFβ/ activin-like SMAD 2/3 signaling (Fig 1C & 1D). These conclusions may help to unify the seemingly incompatible beliefs about the true roles of GDF3 (*4, 9, 11, 16-19*).

**Figure 6:**
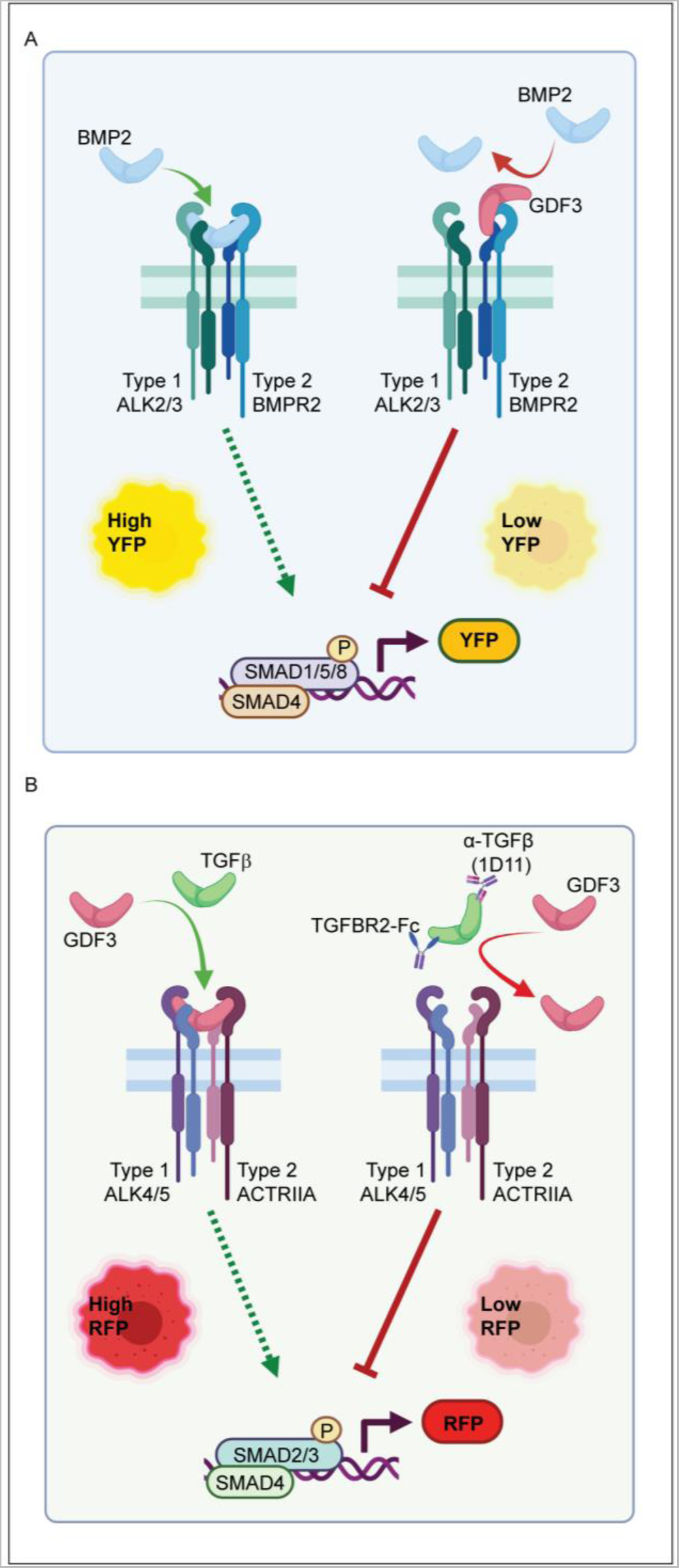
Schematic representation of the role of GDF3 in BMP and Tgfβ signaling. A) GDF3, a member of the Tgfβ superfamily, antagonizes BMP signaling by binding to the BMPR2 receptor. B) GDF3 activates TGFβ signaling either directly by binding to the receptor ACTRIIA or indirectly via direct or indirect interaction with Tgfβ. GDF3’s ability to activate th SBE-RFP repoter is inhibited by TGFβ specific binding agents, anti-TFβ antibody (clone 1D11) or soluble TGFBR2-FC.

### GDF3 is an activin-like BMP signaling antagonist

We present a new mechanism to explain inhibition of BMP signaling by GDF3. The paradigm of GDF3 as a type II receptor antagonist offers an alternative to the model that GDF3 directly interacts with BMP proteins and prevents the formation of BMP homodimers (*4*). Using multiple modalities, we find that GDF3 has a high affinity for the type II receptors BMPR2, ACTRIIA, and ACTRIIB (Fig 3). This promiscuous binding to multiple type II receptors is similar to activin A and activin B, which are also capable of competitive inhibition of BMP signaling (*33*). These effects suggest that the biological role of GDF3 is as an activin-like signaling molecule.

Functionally, our studies provide evidence that GDF3 can broadly inhibit BMP activity. We observe that GDF3 both decreases BMP-stimulated BRE reporter activation and diminishes mRNA expression changes of endogenous BMP2-stimulated genes. We observe these effects both with recombinant mature GDF3 and with the expression of a full-length wild type Gdf3 cDNA. In our experiments, GDF3 was capable of inhibiting reporter activity of all BMPs tested: BMP2, BMP7, BMP9, BMP10, and BMP15. These ligands have varied affinities for the BMPRII, ACTRIIA and ACTRIIB receptors suggesting that GDF3 could inhibit BMP signaling through one or all of these receptors (*33*). Further studies should evaluate which specific type II receptors mediate GDF3’s ability to inhibit different BMP signaling ligands in a cell type dependent manner. Our *in vivo* data also shows increased expression of canonical BMP target genes in a Gdf3 loss of function mouse model. These findings suggest that increased expression of GDF3 seen in obesity may contribute to pathologies linked to diminished BMP signaling. For example, multiple missense mutations in BMPR2 were previously identified in patients with pulmonary arterial hypertension (*34*). Whether GDF3 mimics these effects to affect metabolic function in obesity through BMP signaling antagonism is an area for future investigation.

### GDF3 is a TGFβ/ activin-like SMAD 2/3 signaling agonist

In skeletal muscle, myostatin, activins and TGFβ signaling can inhibit myoblast proliferation and differentiation (*35*). We find that GDF3 shares this biology. RNA sequencing analysis of cells treated with GDF3 suggest an inhibition of cellular differentiation from myoblasts to myotubes (Fig 5D) (*36*). GDF3 is sufficient to activate TGFβ-like reporter activity and expression of TGFβ-induced genes both *in vitro* and *in vivo*. These effects could be mediated by GDF3’s ability to bind to the receptors ACTRIIA and ACTRIIB. These data are in agreement with a previously reported co-immunoprecipitation study of ACTRIIB and GDF3 (*18*). Quite unexpectedly, we found that TGFβ interactors (an anti-TGFβ antibody and soluble TGFBR2-Fc) blocked the ability of either recombinant or genetically encoded GDF3 to activate SMAD 2/3 signaling. However, neither the antibody, nor TGFBR2-Fc bind directly to GDF3. We find instead that GDF3 binds to TGFβ1. Unlike BMPs, GDF3 and activins, TGFβ1-3 only bind to one type II receptor, TGFBR2 (*33, 37*). This suggests that GDF3’s ability to activate SMAD2/3 signaling requires the presence of TGFβ. We propose a model where GDF3 and TGFβ form an active signaling complex. Ligand heterodimerization has been reported for many members of the TGFβ superfamily; whether a GDF3: TGFβ complex is possible and has a physiological role remains to be investigated.

### Physiology: GDF3 in obesity and insulin resistance

In healthy adult animals, *Gdf3* mRNA levels are expressed at very low levels (*38, 39*) but increase with inflammatory conditions including obesity or ischemia (*8, 9, 11, 30, 31*). Mimicking these conditions with acute GDF3 adipose gain-of-function in lean animals promoted a mild insulin resistance with no change in body weight (*9*). Loss of function studies in obese mice provide beneficial phenotypes. Due to the critical role of GDF3 in embryonic development, mice with germline deletion of Gdf3 exhibited decreased survival and developmental abnormalities. While these models are protected from diet-induced obesity, the mechanistic basis for these phenotypes is challenging to interpret (*4, 16, 18, 40*). Limited Gdf3 deficiency was beneficial for glucose homeostasis following Gdf3^-/-^ bone marrow transplantation (*12*). The transplantation model observed reduced activation of the NLRP3 inflammasome and decreased rates of adipose tissue lipolysis. Here we find a complementary phenotype with inducible genetic deletion of Gdf3 in obese adipose tissue leading to improved glucose tolerance and insulin resistance.

BMP and TGFβ play a coordinated role in obesity. Increased BMP signaling enhances thermogenic gene expression in adipose tissue, promoting increased glucose utilization and amelioration of insulin resistance (*41*), while TGFβ1 contributes to tissue fibrosis and inflammation (*42-44*). Our results suggest GDF3 promotes adverse outcomes by promoting gene expression pathways related to TGFβ signaling and to suppress expression of beneficial BMP target genes. As a bi-functional protein, we find that GDF3 loss of function in obesity leads to modestly reduced body weight and improved metabolic health at the physiological level. Using indirect calorimetry, we saw a small but not statistically significant decrease in energy balance that over time may have contributed to the decreased body weight in inducible Gdf3^AKO^ mice. Despite small effects on body weight, we see large differences in gene expression and insulin sensitivity. These findings emphasize the powerful effects of TGFβ superfamily in metabolic physiology.

GDF3 seems to have opposite effects in skeletal muscle or cardiac tissue. Increased GDF3 from infiltrating macrophages or recombinant protein can promote skeletal muscle repair and regeneration after injury (*10, 30, 45-48*). In contrast, cardiac stromal cells secrete GDF3 following ischemic heart injury and promote harmful fibrotic remodeling in the heart (*8*). The gene expression data from our RNA-seq experiments from GDF3 treated cultured myoblasts demonstrates reduced differentiation but also increased TGFβ-like gene expression. Our data can support both models and further highlights the context dependence of gene expression signatures.

### GDF3 gene expression signature

We find BMP target genes upregulated in GDF3^AKO^ mice. These include the Id genes Id1, Id2 and Id4. Id genes are among the first identified genes regulated by BMP signaling and the promoter of Id1 is used for the BRE luciferase and the BRE-YFP reporters. ID proteins belong to a subfamily of helix-loop-helix transcription factors which regulate adipogenesis (*21, 23, 49*). ID1 clearing is essential for adipocyte differentiation. ID2 and ID4 are induced during adipogenesis and are essential for regulating adipose tissue metabolism. ZBTB16 together with HDAC4 plays a major role in matrix mineralization in adipogenic progenitor cells and promotes adipogenesis. Both genes are activated in a BMP6 or BMP2 -dependent manner and regulate the adipose tissue metabolism (*50*). Secretory leukocyte protease inhibitor (SLPI) was previously shown to induce browning in inguinal adipose tissue of mice and thereby lower their body weight by increasing thermogenesis (*51*). In line with these studies, we show that loss of function of Gdf3 in the epididymal adipose tissue enhances the expression of BMP targets and leads to improved metabolic health even on an obesogenic high fat diet.

We find TGFβ targets genes downregulated in GDF3^AKO^ mice. These include Adamts4, Adamts5, and Adamts7, which encode metalloproteinase proteins that degrade the extracellular matrix and are important in tissue remodeling. Also downregulated are Col3a1, Col5a3 and Col18a1 which encode the collagen type III, V and XVIII proteins. These are known TGFβ targets that positively correlate with diet induced obesity and insulin resistance (*52*). The lowered expression of these genes in our Gdf3 loss of function mice could be contributing to the phenotypic effect of improved metabolic health under high fat diet feeding. These gene expression signatures could help to explain the adverse effects of GDF3 after ischemic injury, especially in obesity where GDF3 levels are increased.

GDF3 is an essential factor in embryonic development that is shut down in the postnatal period only to be restored during obesity, inflammation, or ischemia. Expression of TGFβ1 and GDF3 are both increased in obese adipose tissue. Moreover, a SMAD 2/3/4 DNA binding motif was over-represented in H3K27ac promoter and enhancer peaks consistent with increased TGFβ signaling (*44*). GDF3 may be a key modulator of adipose tissue remodeling in obesity. In obesity, a state of “BMP resistance” appears—where despite increasing circulating levels of ligands, BMP signaling is diminished (*53, 54*). Although antagonist ligands noggin and gremlin have been implicated, GDF3 may also be a previously unappreciated component of this biology. Together, these data suggest that GDF3 may play a maladaptive role in the biology of obesity.

## MATERIALS AND METHODS

### Animal models

All animal experiments were performed with approval from the Institutional Animal Care and Use Committees of The Harvard Center for Comparative Medicine, Beth Israel Deaconess Medical Center. Mice were maintained at 12 h/ 12 h light/ dark cycles, 22 ± 2 °C room temperature and 30%-70% humidity with ad libitum access to food and water, in individually ventilated cages. Cages and bedding were changed every two weeks and mice were monitored regularly for their health status by animal technicians with the support of veterinary care and remained free of any adventitious infections for the entire duration of this study. Mice were fed a standard chow diet (13% kcal fat, LabDiet, no. 5053) or a high fat diet (60% kcal fat, Research Diets, no. D12492i) for the indicated durations. Gdf3 floxed mice (Gdf3^fl/fl^) on a C57Bl/6J genetic background were generated using CRISPR/Cas9 with the help of the BIDMC transgenic mouse core. Exon 1 encoding the Gdf3 start codon (ATG) was targeted such that the entire first exon of Gdf3 would be deleted in the presence of Cre recombinase. 8 week old adult male Gdf3^fl/fl^ mice were fed high fat diet *ad libitum* for 8 weeks. They were then anesthetized under isofluorane and bilaterally injected with either AAV2/DJ-CMVeGFP (AAV-GFP) 1.13X10^13^ vg/ml or AAV2/DJ-CMVCre-wtIRESeGFP (AAV-Cre) 1.3x10^13^ vg/ml at a dose of 5ul each into the eWAT. The AAVs were obtained from the University of Iowa Viral Vector Facility, AAV-GFP (VVC-U of Iowa-4382) and AAV-Cre (VVC-U of Iowa-5714). The mice were allowed to recover from surgery for two weeks following which they were subjected to a glucose tolerance test (GTT) 2 weeks after AAV injection and an insulin tolerance test (ITT) 5 weeks post injections. The mice were transferred to indirect calorimetry cages (Promethion, Sable Systems, Inc) 3 weeks post injections to measure their energy expenditure, food and water intake and locomotor activity at room temperature (23°C). Energy balance was calculated from these data using the freely available software CalR version 1.3, developed in house (https://calrapp.org). All mice were euthanized 6 weeks post AAV injections following a 4 hour fast. Tissue samples were harvested and snap frozen in liquid nitrogen before storage at -80°C for further processing.

### Glucose and Insulin Tolerance Tests

Tests were performed on mice fasted for 4 h, with ad libitum access to water. Following an initial blood glucose measurement post-fast, tolerance to an intraperitoneal (IP) injection of glucose (1 g/kg) or insulin (0.75 U/kg) was assessed by change in blood glucose over a period of 120 min. Glycemia was measured by tail vein bleeds at the indicated times using a Contour Next EZ glucometer and glucose strips (Bayer).

### Insulin ELISA

Mice were fasted for 16 hours, 5 weeks post AAV injections and tail vein blood samples were collected in EDTA-coated tubes (Sarstedt, # NC9141704). Plasma was obtained by spinning the blood samples for 10 minutes at 2000g at 4°C. Plasma insulin levels were measured after a 16-hour fast using the Ultra-Sensitive Mouse Insulin ELISA Kit (Crystal Chem, # 90080).

### Plasmid vectors

The pHK3-BRE-Citrine (BRE-YFP) fluorescent reporter as well as the piggyback transposase (p-base plasmid) were gifts from Michael B. Elowitz’s lab (Antebi et al. Cell, 2017). To generate the pHK3-CAGA12-mCherry (SBE-RFP), the BRE-Citrine was replaced with CAGA12-mCherry using the NEBuilder® HiFi DNA Assembly Cloning Kit (New England Biolabs, E5520S). For ease of readability these reporters are referred to as BRE-YFP and SBE-RFP in the manuscript.

The pTetON-TRE-TagBFP2 and pTetON-TRE-mGdf3-P2A-TagBFP2 plasmids were synthesized with codon optimization of mouse Gdf3 cDNA by Vector Builder Inc. GDF3 and/or TagBFP2 (referred to as TetOn-Control and TetOn-Gdf3 in the manuscript) expression were induced using the indicated concentrations of doxycycline for the indicated times. All transfections were carried out using Lipofectamine 3000 (Invitrogen, L3000001) following the manufacturer’s protocol.

### Cell Lines

HEK293T cells (human female in origin) were cultured every 3-4 days using 0.05% Trypsin EDTA and maintained in DMEM (Gibco, 11965118) supplemented with 10% heat inactivated fetal bovine serum (iFBS, Gibco, 10082147) and 1% Penicillin/Streptomycin. C2C12 myoblasts (ECACC, 91031101, C3H female mouse in origin) were cultured every 2-3 days using 0.25% Trypsin-EDTA and maintained in DMEM (GIBCO, 11965118) supplemented with 20% iFBS and 1% Penicillin/Streptomycin. All cell lines were maintained at 37°C at 5% CO_2_.

### Western Blot analysis

HEK293T cells were plated in collagen coated 6-well plates to reach 70-90 % confluency overnight. These cells were then transfected with 2 µg/ well of either the TetOn-Control or TetOn-Gdf3 vectors using Lipofectamine 3000 (Invitrogen) following the manufacturer’s protocol. 24 hours post-transfection, the cells were incubated in serum free media with or without doxycycline for 48 hours. Whole-cell extracts were prepared by washing the cells twice with PBS and then scraping the cells in RIPA lysis buffer (50 mM Tris, pH 7.5, 150 mM NaCl, 1% NP-40, 0.5% sodium deoxycholate, 0.1% SDS) containing 1X HALT protease and phosphatase inhibitor cocktail (Thermo Fisher). Proteins concentrations were quantified using the Pierce BCA protein assay (Thermo Fisher) and protein samples were prepared in reducing Laemmli buffer and heated for 10 minutes at 95°C. Equal amounts of protein were loaded onto precast Mini-PROTEAN TGX gels (Bio-Rad) separated by SDS-PAGE using the appropriate Bio-Rad apparatus and transferred to PVDF membranes using the Trans-Blot Turbo transfer system (Bio-Rad). The blots were blocked with 5% BSA in TBST and blotted according to manufacturer’s recommendations for the indicated primary and secondary antibodies (Table S2) followed by development with the femtoLUCENT™ PLUS-HRP Chemiluminescent kit (G-Biosciences) for the anti-GDF3 blot and the SuperSignal™ West Pico Chemiluminescent kit (Thermo Fisher) for anti-β-Actin blot and the BioRad Chemidoc Touch Imaging system.

### Generation of dual reporter cells

C2C12 myoblasts were transfected with the BRE-YFP reporter plasmid and the piggyback transposase (p-base plasmid) designed to be stably integrated using the piggybac transposon system (*55*). Stable cells were generated by selection with 800µg/mL of Hygromycin B (Sigma-Aldrich, H7772). The cells were again transfected with the SBE-RFP reporter with the piggyback transposase and selected for stable integration of both reporters by three rounds of fluorescence-assisted cell sorting for YFP and RFP double positive cells (dual reporter cells).

### Flow cytometry

C2C12 dual reporter cells were plated in a 96 well cell culture plate to reach 50 to 70% confluency the next day in complete medium (DMEM high glucose with 20% heat inactivated fetal bovine serum (iFBS) and penicillin and streptomycin. Cells were then incubated with the specified concentrations of recombinant proteins in complete medium. If cells needed to be serum starved, the medium was replaced with serum starvation medium (DMEM high glucose with 0.2-0.3% fatty acid free bovine serum albumin and penicillin and streptomycin) for 4 hours the day after plating followed by incubation with recombinant proteins in serum-free media for 24 hours.

For the induction of genetically encoded Gdf3 and/or BFP, dual reporter cells were transiently transfected with either the TetOn-Control or the TetOn-Gdf3 plasmids in 10 cm cell culture dishes. After an overnight incubation in complete medium, these cells were split into a 96 well cell culture plate. After another overnight incubation in complete medium, these cells were serum starved for 4 hours and then treated with doxycycline in serum-free media for the indicated time points. The cells were then processed for flow cytometry.

Briefly, the medium from the 96 well plate was removed and cells were washed with PBS. Cells were then dissociated with 0.25% Trypsin EDTA in a cell culture incubator for approximately 5 minutes. Trypsin was neutralized with PBS containing 3% iFBS and followed by spinning the plate in a refrigerated centrifuge at 500g for 10 minutes with a swinging rotor. The supernatant was dumped and FACS buffer (PBS + 0.3% BSA) containing 1:10000 LIVE/DEAD Fixable Far Red Dead Cell Stain (Thermo Fisher Scientific, #L34973) was added to the cells for 15 minutes on ice. The plate was then centrifuged again, the medium was removed and the plate was washed with FACS buffer and centrifuged again. The supernatant was removed again and each well was incubated with 200uL of FACS buffer. These cells were then analyzed using the CytoFLEX flow Cytometer (Beckman Coulter) and the CytExpert acquisition and analysis software. Briefly, single unstained C2C12 cells and single fluorescent controls were used to set the gains and gates for each fluorescent channel. Cells were gated to analyze 1000 live single cells from each well. The mean fluorescence intensity (MFI) of either YFP or RFP of the live cells from each well was used for the subsequent analysis. For the experiments with doxycycline induced BFP or GDF3 and BFP expression, the MFI of YFP or RFP was measured from cells expressing YFP, RFP and BFP.

### RNA sequencing

Dual reporter cells were plated in 12 well plates to reach 70% confluence overnight. The cells were then incubated in serum starvation media for 4 hours followed by incubations with recombinant proteins in serum starvation media for 24 hours. The cells were then washed with PBS and lysed in Tri-reagent and RNA was extracted using the Direct-zol RNA miniprep kit (Zymo Research, # R2050) following the manufacturer’s instructions. Total RNA was quantified using the NanoDrop 2000 spectrophotometer (Thermo Fisher Scientific, # ND-2000). mRNA was enriched from 400ng of total RNA using the NEBNext rRNA Depletion Kit v2 (NEB, # E7400X) to remove ribosomal RNA. First stand was synthesized using the Random Hexamer Primer (Thermo Fisher Scientific, # SO142) and Maxima Reverse Transcriptase (Thermo Fisher Scientific, #EP0742). Second strand was synthesized using NEBNext mRNA Second Strand Synthesis Module (NEB, #E6111L). Sequencing adaptors were added by tagmentation and amplified for 12 cycles using the Nextera XT DNA Library Preparation Kit (Illumina, # FC-131). Libraries were pooled at a final concentration of 1.35 pM, and 36bp x 36bp paired end reads were sequenced on an Illumina NextSeq500 instrument. Sequencing reads were demultiplexed using bcl2fastq and aligned to the mm10 mouse genome using HISAT2 (a fast spliced aligner with low memory requirements) (*56*). PCR duplicates and low-quality reads were removed by Picard (https://broadinstitute.github.io/picard) and filtered reads were assigned to the GRCm38 genome modified to minimize overlapping transcripts and quantified using featureCounts (an efficient general purpose program for assigning sequence reads to genomic features (*57*). Differential expression analysis of the data was performed using EdgeR (*58*).

Analysis of gene sets regulated by Gdf3 was performed with Enrichr. This resource introduced us to Sysmyo, a large collection of signatures of gene expression changes in muscle. (*36, 59*).

### RTqPCR

Dual reporter cells were plated in 6 well plates and treated as described in the figure legends. Cells were lysed in Tri-reagent and RNA was extracted using the Direct-zol RNA miniprep kit (Zymo Research, # R2050) following the manufacturer’s instructions. Quantitative real-time PCR (qPCR) was performed using cDNA generated with the High Capacity cDNA Reverse Transcription kit (ThermoFisher Scientific, # 4368813) with SYBR Select Master Mix (Applied Biosystems, # 4472920) on the QuantStudio 6 Real-Time PCR System (Applied Biosystems). Relative mRNA expression was determined by the 2^-ddCt method normalized to TATA-binding protein (TBP) levels. The sequences of primers used in this study are listed in Table S3.

### Phylogenetic analysis

Phylogenetic analysis of human TGFβ superfamily proteins was performed with ClustalW (*60*). All sequences were retrieved from UniProt (uniprot.org) using the database’s annotation for the mature chain. Alignments were performed with the blosum62mt2 matrix, ignoring positions with gaps. Human GDF3 was used as the starting profile for alignment.

### Surface Plasmon Resonance (SPR)

SPR was performed in 20 mM HEPES pH 7.4, 350 mM NaCl, 3.4 mM EDTA, 0.05% P-20 surfactant, 0.5 mg/ml BSA at 25℃ on a Biacore T200 optical biosensor system (Cytiva). Human Fc-fusion constructs of each receptor were purchased from R&D Systems (TGFβRII-Fc: Cat. No. 341-BR-050/CF, BMPRII-Fc Cat. No. 811-BR-100/CF, ActRIIA-Fc Cat. No. 340-R2-100/CF,

ActRIIB-Fc Cat. No. 339-RB-100/CF). Each receptor was captured using a series S Protein A sensor chip (Cytiva) and immobilized with a target capture level of ∼70 RU with a contact time of 60 seconds at 20 µL/min flow rate. A 16-point, two-fold serial dilutions were performed for rhGDF3 (R&D systems, Cat. No. 5754-G3-010/CF) at a starting concentration of 12.5 nM to 0.391 nM. Each cycle had an association of 200 seconds and disassociation of 900 seconds at a flow rate of 50 µL/min. The sensor chip was regenerated with 10 mM Glycine pH 1.7. Kinetic analysis was utilized on the Biacore T200 evaluation software and fit by 1:1 fit model with mass transport limitations. All SPR experiments were performed in triplicate, except TGFβRII-Fc which was run in duplicate and where no binding was observed. Curves were fit individually using a 1:1 model for kinetic binding using Biacore software and plotted with GraphPad Prism. The list of recombinant proteins used are listed in Table S1

### Biolayer Interferometry (BLI)

All binding assays were performed with the Gator Plus (Gator Bio) instrument using 96-tilted well plates. Anti-Human IgG Fc Gen II (#160024) and Anti-Mouse IgG Fc (#PL168-160004) probes, BLI 96-Flat Plate Polypropylene (#130150), Max Plate (#130062), and Regen Buffer no salt (#120063) were all purchased from Gator Bio. Samples were diluted in freshly prepared and filtered running buffer containing 1x phosphate buffered saline, 1 mg/mL BSA, and 0.1% Tween20, pH 7.4 at 25°C with an orbital shake speed of 1000 rpm. Kinetic assays were performed by first equilibrating the sensors for 600 seconds in kinetic buffer, followed by 150 seconds of baseline, 200 seconds of loading, 150 seconds of washing, 300 seconds of association, 300 seconds of dissociation, and 5 cycles of regeneration (5 seconds regeneration/5 seconds neutralization). All the capturing receptors and antibodies were used at a final concentration of 10 µg/mL. The target proteins were used at different concentrations as described in the results section and the respective figure legends. The data were analysed using the proprietary software offered by Gator Bio®. The list of recombinants and antibodies used are listed in Table S1 and S2.

### Isothermal Titration Calorimetry (ITC)

The incremental ITC binding experiments were performed in the Nano ITC Standard Volume with a fixed cell by titrating 250ng/mL of GDF3, 250 ng/mL or 2.5ng/mL of TGFβ, and 250ng/mL of BMPR2-Fc into 1µg/ml of 1D11 or 250ng/mL of GDF3. The incremental ITC experiments consisted of 10, 8μL injections at 350-second intervals with stirring speed of 300 revolutions per minute (rpm). The data was acquired using ITC Run data acquisition software. ITC data were analyzed with Nano Analyze Software. The list of recombinant proteins and antibodies used are listed in Table S1 and S2.

### Statistical Analyses

Data were analyzed using Prism software (GraphPad Software) and are expressed as mean ± SEM. All data were assumed to have a normal distribution and two-tailed Student’s t tests, one-way ANOVA with Newman-Keuls Multiple Comparison test, or two-way ANOVA were used to compare means between groups as indicated in the figure legends; p < 0.05 was considered significant.

### Data Availability

At the time of this submission, we are in the process of submitting the RNA-seq dataset to the NIH SRA.

## ACKNOWLEDGEMENTS

Financial support for this project was provided to A.S.B. by NIH (R01DK107717, R01DK133948, S10OD028635). A.S.B. has also received support for an investigator-initiated grant from Eli Lilly & Co. RNA-seq was performed by the Boston Nutrition and Obesity Research Center’s Functional Genomics and Bioinformatics Core (BNORC, NIH 5P30DK046200). We thank Christopher Jacob from the Functional Genomics and Bioinformatics core at BIDMC for his help with RNA seq data analysis. We also thank Michael B. Elowitz from Caltech for the BRE-Citrine vector. We thank Viet Le from Boston Children’s Hospital for helpful discussions for the gift of recombinant TGFβ1 and activin A. The authors have no other relevant competing interests to declare.

## SUPPLEMENTARY MATERIAL

**Figure S1.**
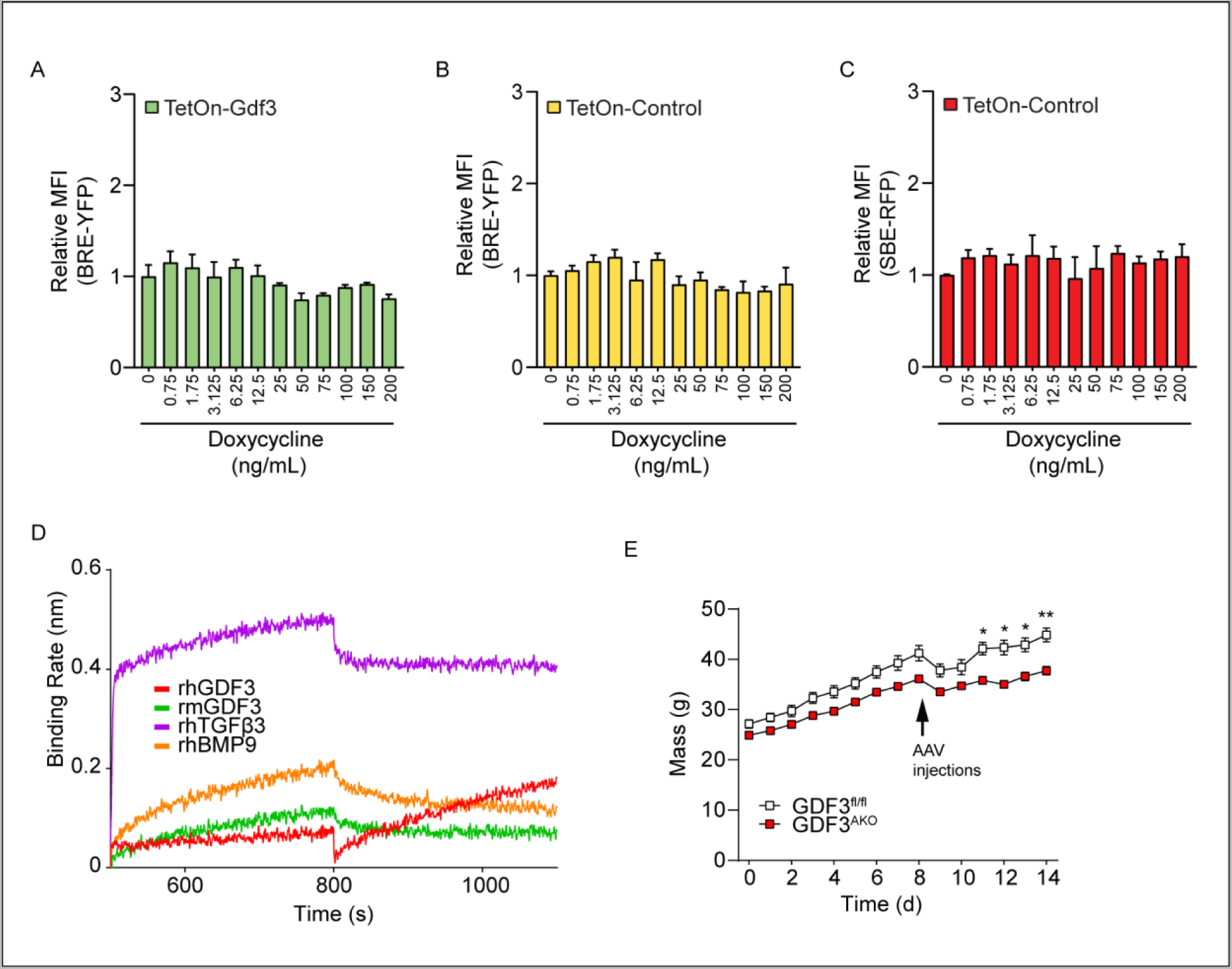
A-C) C2C12 dual reporter cells were transiently transfected with TetOn-Gdf3 or TetOn-Control vectors for 24 hours followed by doxycycline treatment in serum free media for 48 hours. Cells were analyzed by flow cytometry for changes in YFP reporter activitiy (A, B) or RFP reporter activity (C). D) BLI analysis of 100nM each of rhGDF3, rmGDF3, rhTGFβ3 or rhBMPƏ with 10 µg/mL of anti-TGFβ antibody (clone 1D11). E) Body weights of GDF3^fl’fl^ and GDF3^AK0^ male mice from beginning of HFD feeding (week 0). *p<0.05, **p<0.01, ***p<0.001, ****p<0.0001 compared to control (Gdf3^fl/fl^).

**Table S1.** Recombinant proteins.

| Recombinants | Species | Catalog No | Company | Used for |
| --- | --- | --- | --- | --- |
| GDF3 | Mouse | 9009-GD-010 | R&D Systems | Flow Cytometry, BLI ITC |
| GDF3 | Human | 5754-G3-010 | R&D Systems | BLI |
| GDF3 | Human | 5754-G3-010/CF | R&D Systems | SPR |
| BMP2 | Human | N/A | Vicki Rosen lab | Flow Cytometry |
| BMP7 | Mouse | 5666-BP-010 | R&D Systems | Flow Cytometry |
| BMP9 | Human | 3209-BP-010 | R&D Systems | Flow Cytometry |
| BMP10 | Mouse | 6038-BP-025 | R&D Systems | Flow Cytometry |
| BMP15 | Human | 5096-BM-005 | R&D Systems | Flow Cytometry |
| TGFβ1 | Human | T7039 | Millipore Sigma | Flow Cytometry, ITC |
| TGFβ3 | Human | 8420-B3-025 | R&D Systems | BLI |
| Activin A | Human/ Mouse/ Rat | 338-AC-010 | R&D Systems | Flow Cytometry |
| BMPRII-Fc | Human/ Mouse | 811-BR-100 | R&D Systems | SPR, BLI, ITC |
| TGFBRI-Fc | Human | 3025-BR-050 | R&D Systems | Flow Cytometry |
| TGFBRII-Fc | Human | 341-BR-050 | R&D Systems | Flow Cytometry |
| TGFBRII-Fc | Human | 341-BR-050/CF | R&D Systems | SPR |
| TGFBRII-Fc | Human | 241-R2-025 | R&D Systems | BLI |
| TGFBRIII-Fc | Mouse | 5034-R3-050 | R&D Systems | Flow Cytometry |
| ACTRIIA-Fc | Human | 340-R2-100/CF | R&D Systems | SPR |
| ACTRIIA-Fc | Human | 340-RC2-100 | R&D Systems | BLI |
| ACTRIIB-Fc | Human | 339-RB-100/CF | R&D Systems | SPR |
| ACTRIIB-Fc | Human | 339-RB-100 | R&D Systems | BLI |

**Table S2.**
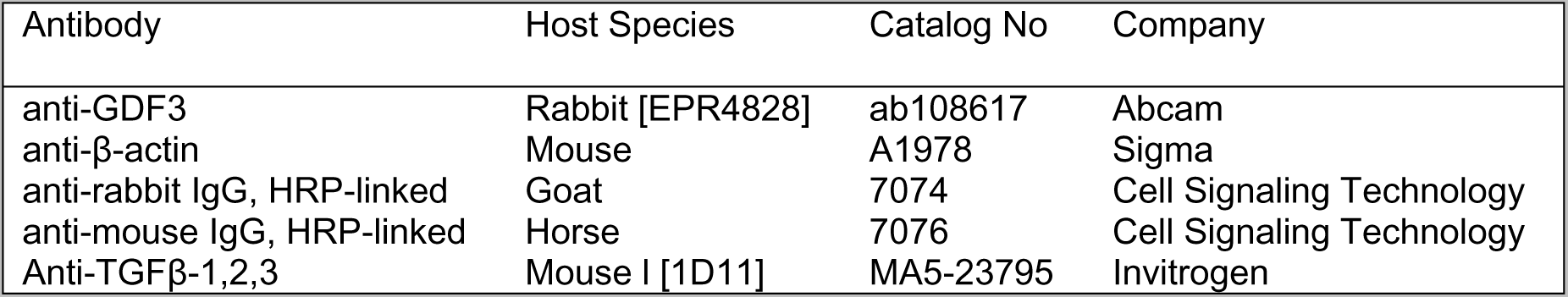
Antibodies.

**Table S3:**
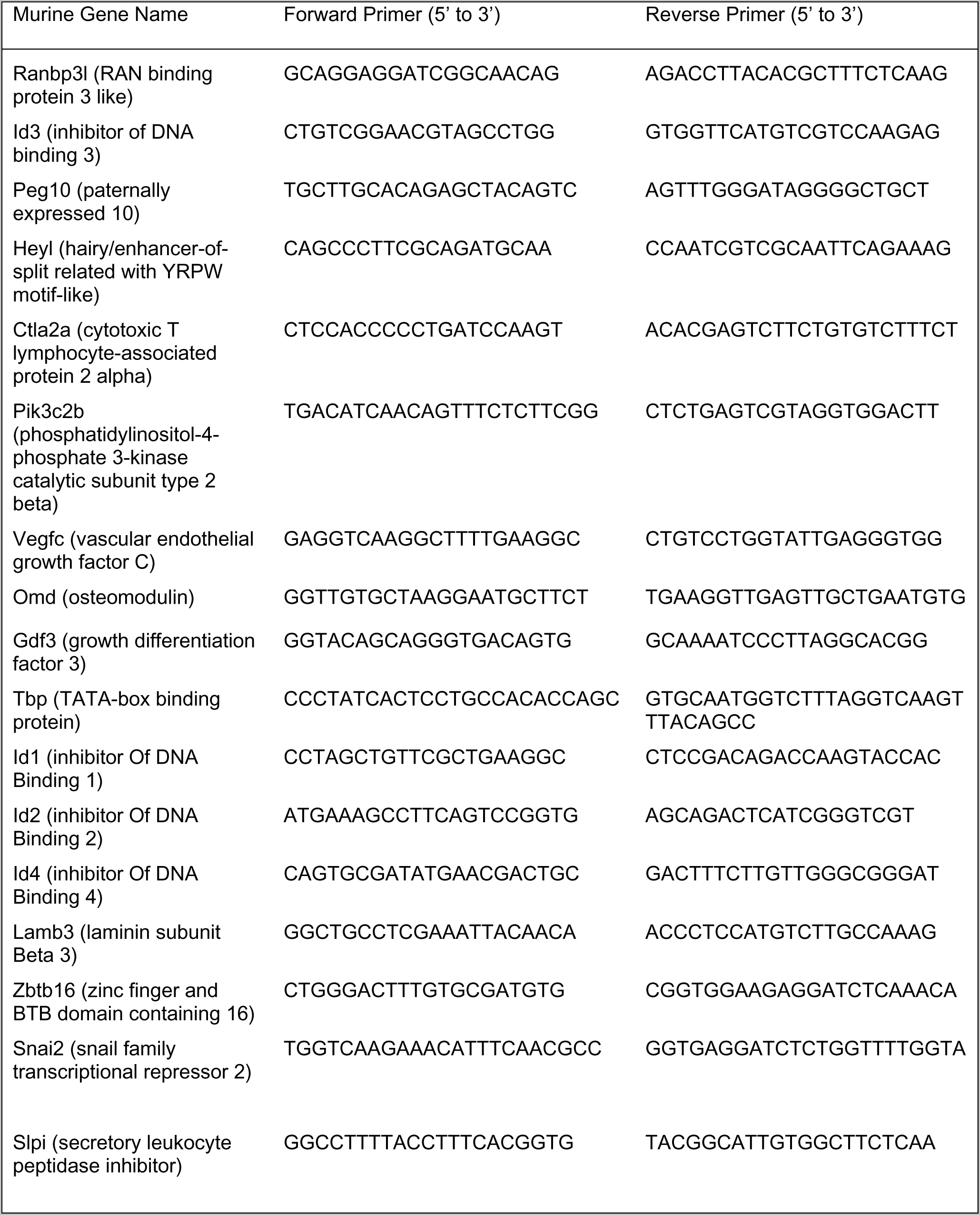
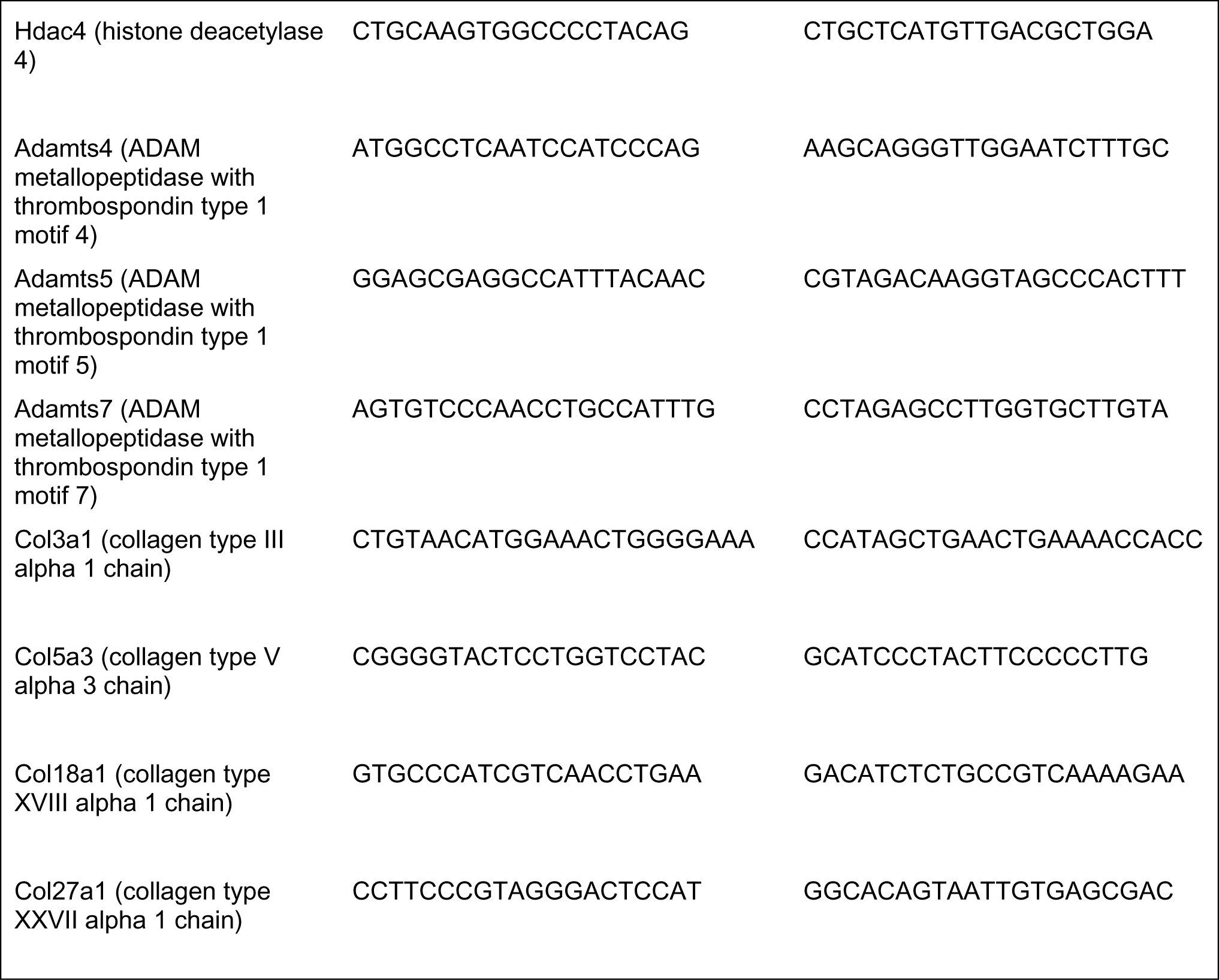
qPCR primer list.

